# Thermosensitivity of TREK K2P channels is controlled by a PKA switch and depends on the microtubular network

**DOI:** 10.1101/2024.11.07.622439

**Authors:** Sönke Cordeiro, Marianne Musinszki

## Abstract

Temperature sensing is an essential component of animal perception and enables individuals to avoid painful or lethal temperatures. Many temperature sensors in central and peripheral neurons are ion channels. Here, we focus on the thermosensitive TREK/TRAAK subfamily of K2P channels – the only known K^+^ selective thermosensitive channels. The C-terminal domain is essential for the temperature activation of TREK channels, but the mechanism of temperature sensation and the nature of the temperature sensor are unknown. We studied the thermosensitivity of representatives of all K2P channel subfamilies and identified TREK-1 and TREK-2 as the only thermosensitive K2P channels, while TRAAK, the third member of the mechano-gated subfamily, showed no temperature dependence. We transferred the thermosensitivity of TREK-1 to TRAAK channels by exchanging the C-termini, demonstrating that the C-terminal domain is sufficient to confer thermosensitivity. By gradually truncating the C-terminus, we isolated a specific temperature responsive element (TRE) consisting of 18 amino acids that constitutes a unique feature in mammalian thermosensitive channels. Within this TRE lie both the binding domain for microtubule associated protein 2 (MAP2) and the PKA phosphorylation site. Pharmacological disruption of the microtubular network as well as the loss of the MAP2 binding site suppressed the temperature response, and PKA activation completely abolished temperature sensitivity. Thus, the connection to the microtubular network enables the thermosensitivity of TREK channels, which is not intrinsic to the channel itself, while the PKA-mediated phosphorylation status acts as a switch that determines if TREK channels are thermosensitive at all.

## Introduction

Sensing temperature is of vital importance for homoiothermic animals. For the maintenance of a constant body temperature it is sensed mainly by thermosensitive neurons in the preoptic area (POA), a part of the hypothalamus (Morrison, Nakamura and Madden, 2008). To maintain body temperature upon changes in environmental temperature and to avoid harmful temperatures, thermosensitive neurons are located in dorsal root ganglia (DRG) and trigeminal ganglia (TG) (Palkar, Lippoldt and McKemy, 2015), where different ion channels mediate the thermosensation. A measure for temperature sensitivity of biological processes is the change of the rate of a process upon changing the temperature by 10 °C, expressed as the Q10 value. Every enzyme or ion channel has Q10 values up to 3 (Hille, 2001; Sengupta and Garrity, 2013). Only very few ion channels are in fact temperature-sensitive, with Q10 values sometimes higher than 20. Major players are members of the transient receptor potential (TRP) channels, which are intensively investigated, as e.g. disruption of the gene encoding TRPV1 leads to a strong impairment of noxious heat sensing (Caterina *et al*., 2000; Davis *et al*., 2000), while knockout of its antipode TRPM8 causes a suppression of cold sensation (Bautista *et al*., 2007; Colburn *et al*., 2007; Dhaka *et al*., 2007). Several other TRP channels are activated by temperature changes (TRPV2, TRPV3, TRPV4, TRPM3, TRPA1 and TRPC5; Palkar, Lippoldt and McKemy, 2015), but despite an intensive search, no distinct thermosensor could yet be identified in these channels.

In addition to ‘classical’ temperature sensitive TRP channels, members of other ion channel families have also been shown to be activated by temperature changes: the voltage-dependent Na^+^ channel Na_v_1.8 (Zimmermann *et al*., 2007), the Ca^2+^ activated Cl^-^ channel TMEM16A (also known as anoctamin 1; Cho *et al*., 2012), and the STIM1/Orai pair (Xiao *et al*., 2011). The members of the TREK/TRAAK subfamily of K2P channels are the only known thermosensitive K^+^-selective channels. Importantly, in contrast to the thermosensitive ion channels described above, activation of these K^+^ channels dampens neuronal activity (Lamas, Rueda-Ruzafa and Herrera-Pérez, 2019).

While all other K^+^ channels are composed of four pore forming subunits and subsequently possess a tetrameric symmetry, K2P channels are composed of only two subunits with four transmembrane domains and two pore segments in tandem. According to their homology and similarity in activation mechanisms, the 15 members of the mammalian K2P channel family are subdivided into six subgroups – the weak inward rectifiers (TWIK), the mechano-gated (TREK/TRAAK), the alkaline-activated (TALK), the Ca^2+^-activated (TRESK), the acid-inhibited (TASK), and the halothane-inhibited (THIK) subfamily (Honoré, 2007; Enyedi and Czirják, 2010).

All members of the TREK/TRAAK subfamily are expressed in thermosensitive neurons of the POA and the DRG and TG (Maingret *et al*., 2000; Talley *et al*., 2001; Yamamoto, Hatakeyama and Taniguchi, 2009; Marsh *et al*., 2012; Viatchenko-Karpinski, Ling and Gu, 2018). TREK/TRAAK channels are of essential meaning for the temperature-dependent behavior in mammals. Knockout of one or more of these channels in rodents results in significantly changed heat sensation (Noël *et al*., 2009; Pereira *et al*., 2014; Lamas, Rueda-Ruzafa and Herrera-Pérez, 2019).

TREK/TRAAK channels are regulated by many physical and cellular stimuli, i.e. temperature, mechanical stress, intracellular pH, lipid species like arachidonic acid and PIP2 as well as G protein coupled receptor (GPCR)-mediated phosphorylation and protein-protein interactions (Enyedi and Czirják, 2010). Significantly, truncated TREK-1 channels lacking the intracellular C-termini have reduced activity at rest (Patel *et al*., 1998), and show impaired responses to said stimuli: reduced temperature activation (Maingret *et al*., 2000), reduced mechano-sensitivity (Patel *et al*., 1998), reduced sensitivity to acidification of the intracellular milieu (Maingret *et al*., 1999), and to lipids like arachidonic acid and PIP_2_ (Chemin *et al*., 2005). Thus, for the action of most physiological modulators, the C-terminus is essential (Enyedi and Czirják, 2010; Bagriantsev, Clark and Minor, 2012). However, the mechanism how temperature may be sensed by the C-terminal domain (CTD) to open the channel remains unclear.

Here, we investigate the mechanism of temperature-induced activation of TREK-1 K2P channels and we identify the temperature responsive element (TRE) necessary for their temperature activation. We demonstrate that thermosensitivity is not an intrinsic property of TREK-1 channels, but requires contact to the cytoskeleton that is mediated by microtubular adaptor proteins. Finally, we describe that thermosensitivity is controlled by the PKA-mediated phosphorylation status of the channel, enabling the cell to tune thermosensitivity according to its current state.

## Material and methods

### Molecular biology

Channel constructs were subcloned into the expression vector pFAW containing a CMV promotor. All K2P channels correspond to the human isoforms TREK-1 (KCNK2; NM_001017425), TREK-2 (KCNK10; NM_021161), TRAAK (KCNK4; NM_033310), TALK-2 (KCNK17; NM_031460), TRESK (KCNK18; NM_181840), TWIK-1 (KCNK1, NM_002245), TASK-3 (KCNK9, NM_001282534) and THIK-1 (KCNK13, NM_022054). For control measurements the mouse TRAAK was used (KCNK4; NM_008431). For receptor coupled modulation of temperature - sensitivity, the mouse dopamine receptor D1 (Drd1, NM_001291801) was coexpressed with the TREK-1 channel. For measurements of homomeric TWIK-1 channels, the endo-/lysosomal retrieval motif was destroyed by substituting isoleucine 293 and 294 for alanine. Amino acid substitutions were introduced via site-directed mutagenesis with the QuikChange method (Stratagene). Channels were truncated by replacing the respective amino acid triplet by a stop codon and appropriate restriction site. Chimeric channels were constructed by the insertion of MluI restriction sites in TREK-1, TRAAK, TALK-2 and TASK-3 by silent mutation in L311/R312/V313 (TREK-1), L257/R258/V259 (TRAAK) and G271/R272/V273 (TALK-2) and by replacing Phe246 (TASK-3). Consequently, chimeras consist of TREK-1 (M1-V313)/TRAAK (V260-V393), TRAAK (M1-V259)/TREK-1 (I314-K426), TALK-2 (M1-V273)/TREK-1 (I314-K426), and TASK-3 (M1-A245)/TREK-1 (V313-K426). Heteromeric channels were constructed by concatenating TRAAK and TREK-1 with a linker sequence consisting of a XhoI restriction site followed by 10 amino acids (GGGGSGGGGS).

### Cell culture

HEK293 cells were kept in Dulbecco’s Modified Eagles Medium (DMEM) supplemented with 10 % FCS and penicillin-streptomycin (100 U ml^-1^/100 µg ml^-1^) in a 5 % CO_2_ incubator at 37 °C. The cells were transiently transfected with Lipofectamine 2000 (Invitrogen). For electrophysiological measurements, the transfected cells were trypsinized and seeded onto 10 mm coverslips at least 4 h before the experiments.

### Electrophysiology

Most electrophysiological measurements were done in the whole-cell configuration of the patch-clamp technique using an EPC10 amplifier (HEKA) and the PatchMaster software (HEKA). The cells were stimulated by a ramp protocol between -100 and +60 mV (1 s duration, every 5 s, holding potential -80 mV) or by a family of rectangle pulses (1 s duration, between -80 and +60 mV with 20 mV increments, holding potential -80 mV). Pipette resistances were 1-3 MΩ when filled with intracellular solution (in mM): 140 KCl, 2 MgCl_2_, 1 CaCl_2_, 2.5 EGTA, 10 HEPES, pH 7.3 with KOH. For measurements of TWIK-1 channels, K^+^ was replaced by Rb^+^ as TWIK-1 channels display nearly no K^+^ conductance. Where needed, 3 mM MgATP and 0.3 mM NaGTP were included in the intracellular solution. The bath solution contained (in mM): 135 NaCl, 5 KCl, 2 MgCl_2_, 2 CaCl_2_, 10 glucose, 10 HEPES, pH 7.3 with NaOH. On-cell and inside-out measurements were performed with the following bath solution (mM): 140 KCl, 2 MgCl_2_, 2 CaCl_2_, 10 glucose, 10 HEPES, pH7.3 with KOH. All modulatory agents were added to the bath to obtain the specified final concentrations. The temperature was changed via a water-heated and -cooled bath chamber connected to a peltier element and controlled by a temperature controller (TEM-01D, npi electronics). Fast temperature changes were obtained by heating with a halogen lamp (Osram, 15 V, 150 W) mounted in direct vicinity of the recording chamber, yielding heating rates between 0.04 and 0.5 °C s^-1^.

### Chemicals

Forskolin, Phorbol-12-Myristate-13-Acetate, Vinblastine and H-89 were purchased from Cayman Chemical, and MgATP, NaGTP, GMP PNP, 3-Isobutyl-1-Methylxanthine, Colchicine, Cytochalasine B, arachidonic acid and BL-1249 from Sigma-Aldrich. GTPγS was purchased from Jena Bioscience.

### Data analysis

All data are given as mean ± S.E.M. The fold change of temperature activation was estimated by dividing the maximally activated current by the basal current immediately before temperature elevation. Q10 values were calculated by the following equation: 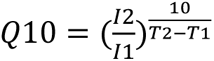 where T1 and T2 are temperatures at different time points and I1 and I2 are the respective currents measured at 0 mV. For intra-individual data comparisons, the Wilcoxon-rank-test was performed using the Igor Pro software (WaveMetrics). Differences were considered significant if p<0.05 and values of p<0.05, p<0.01, and p<0.001 are depicted by *, **, and ***, respectively. All results are summarized in Table S1.

## Results

### Only TREK-1 and TREK-2 are thermosensitive K2P channels

We systematically investigated the thermosensitivity of members of all six K2P channel subfamilies, with a focus on the TREK/TRAAK subfamily for which temperature sensitivity is reported. We measured the current response of channels expressed in transiently transfected in HEK293 cells upon heating, fast heating, or cooling of the bath solution. Both TREK-1 and TREK-2 channels displayed strong activation by increased temperature, while representatives of other subfamilies were not activated (Fig. 1A). The time course of temperature dependent activation is shown in Fig. S1A for a selection of K2P channels. Q10 values for the activation of TREK-1 and TREK-2 channels reached >20 and >15, respectively, but varied strongly depending on the temperature and on the speed of heating (Fig. S1B). Therefore, instead of using the Q10 values, we determined the fold change of the maximal temperature-activated current across the whole temperature range.

**Figure 1.**
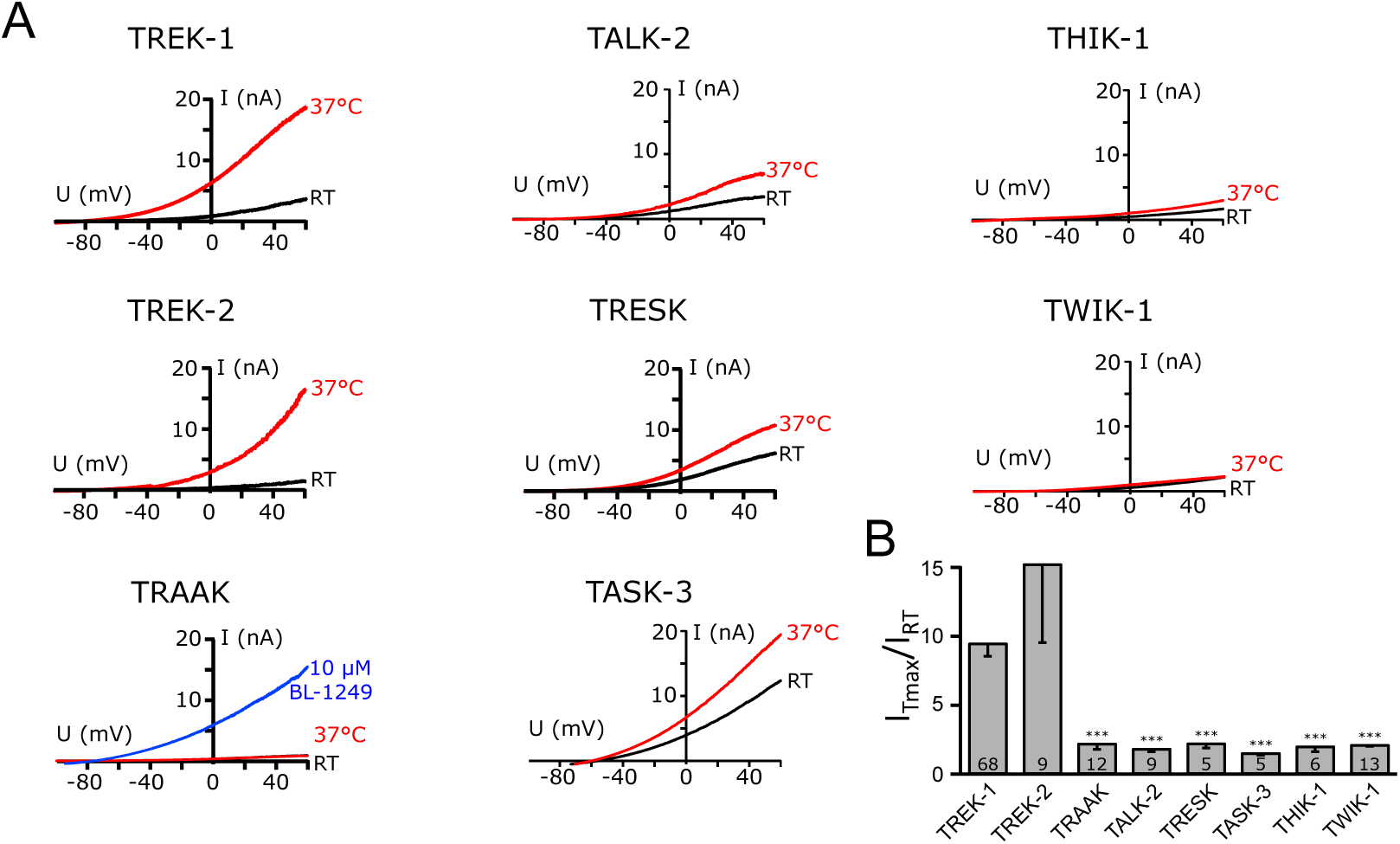
Temperature sensitivity of the TREK/TRAAK subfamily and representative members of all other K2P subfamilies. **A** Exemplary whole-cell current traces recorded from HEK293 cells transfected with the K2P channels at RT (black) and 37 °C (red). Currents were elicited with ramp protocols between -100 and +60 mV. As TRAAK channels showed no currents in the whole cell configuration, these channels were activated by 10 µM BL-1249 after temperature elevation. **B** Fold change of temperature activation of the tested K2P channels (as the maximal current measured at +60 mV normalized to basal current before temperature activation).

TREK-1 and TREK-2 channels displayed a clear temperature-dependent activation with a fold change of the current of 10.53 ± 1.24 and 13.67 ± 5.25, respectively. Elevating temperature did not activate any other tested K2P channel. They showed fold changes of 1.88 ± 0.25 for TRAAK, 1.88 ± 0.21 for TALK-2, 2.25 ± 0.36 for TRESK, 1.54 ± 0.21 for TASK-3, 2.03 ± 0.41 for THIK-1, and 2.25 ± 0.19 for TWIK-1 (Fig. 1B). Surprisingly, the third member of the TREK/TRAAK subfamily, TRAAK, was not temperature sensitive. The functionality of the TRAAK channels was tested by the subsequent activation of the channels by a known pharmacological TREK/TRAAK channel activator (10 µM BL-1249). This observation conflicts with a previous study by (Kang, Choe and Kim, 2005), who reported that TRAAK channels are activated by elevated temperature. However, the study used murine TRAAK, which differs from the human homolog especially in the sequence of the CTD (Fig. S2A). To rule out a species-dependent effect, we also measured the temperature activation of mouse TRAAK channels. Consistent with our previous results in human TRAAK, no temperature-dependent activation was present and the fold change of 3.44 ± 0.29 was clearly below the corresponding values for TREK-1 and TREK-2 channels (Fig. S2B, C). To exclude a possible influence of the cellular background, we also determined the temperature sensitivity of human TRAAK channels expressed in COS7 cells, and again found no activation (fold change 2.07 ± 0.33; Fig. S2C).

It is worth noting that within the representatives of other K2P subfamilies, only TWIK-1 mediated currents also responded to elevated temperature. However, instead of being activated, TWIK-1 channels became inactivating upon depolarization with continuous heating to 37°C (Fig. S3), leading to a current decrease below the initial current at room temperature. This phenomenon has also been described for TWIK-2 channels (Patel *et al*., 2000).

Thus, we find that TREK-1 and TREK-2 channels are the only K2P channels that are activated by elevated temperatures.

### Temperature sensitivity is not intrinsic to TREK-1 channels

The temperature-induced current increase over time disclosed two features: TREK channels activated slower than the temperature rise, and currents decreased upon further holding of the temperature in the whole-cell configuration (Fig. 2A). The delays in time courses of current activation were nearly independent from the heating speed, reaching their maxima at the earliest one minute after starting the temperature rise (Fig. S1A, S4A; compare time courses with a heating rate of 0.1 °C s^-1^ and 0.25 °C s^-1^). Consequently, also the temperatures at maximal current depended significantly on the heating speed (Fig. S4B; note maximal currents at 36 °C and 49 °C). TREK-1 channels are also sensitive to cooling below room temperature, which results in current decrease (Fig. 2D). Cooling of the cells led to a nearly complete closure of TREK-1 channels, and subsequent heating activated the channels as usual with a maximum at 40 °C (Fig. S4D). With this temperature protocol (i.e. cooling to 10 °C before heating) the maximal current was also reached after about two minutes, showing that no specific threshold for temperature activation exists within the applied temperature range.

**Figure 2.**
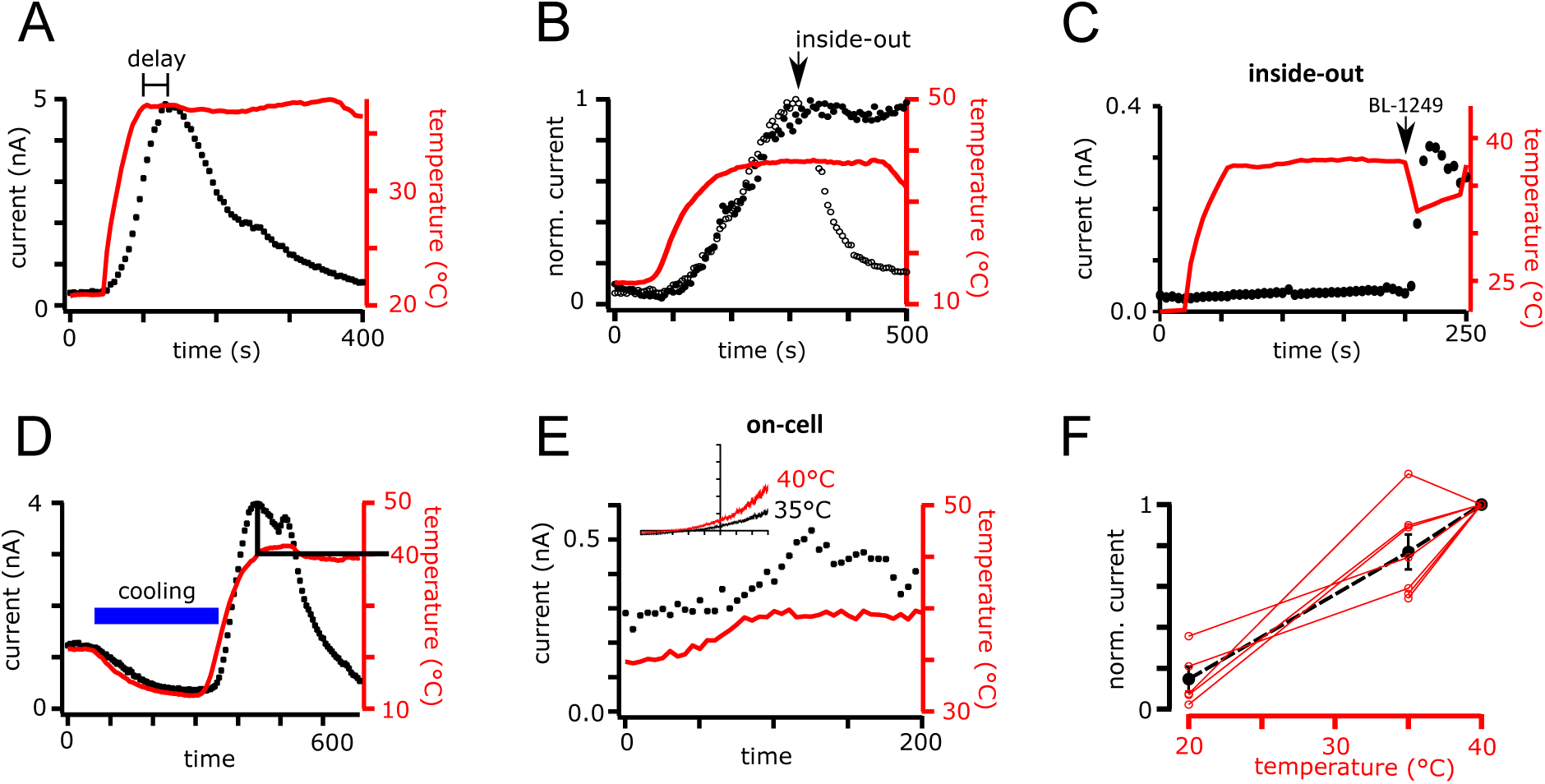
Temperature response of TREK-1 channels depends on the recording mode during activation. **A** Time course of temperature activation in whole-cell recordings of TREK-1 channels. Upon fast temperature elevation, a delay between the maxima of temperature and current is visible. Currents were elicited with a ramp protocol (-100 to +60 mV) and currents measured at +60 mV were plotted over time. **B** Comparison of the time course of temperature activation in the on-cell mode (filled circles) with the time course in inside-out configuration, showing the inactivation upon patch excision after reaching maximal activation (open circles; arrow marks moment of excision). Currents were measured with the ramp protocol in symmetrical K^+^ solutions. **C** Time course of currents measured in the inside-out mode with temperature elevation. As no currents were activated, the channels were activated by 10 µM BL-1249 after temperature elevation as a control (arrow). **D** Time course of temperature activation measured as in A but including a cooling step to ∼10 °C before temperature elevation. **E** On-cell recording of temperature activation in the physiological relevant temperature range (35 °C to 40 °C). Currents were measured as in B. The inset shows exemplary ramps measured at 35 °C (black) and 40 °C (red) from the same patch. **F** Current response of measurements like in E plotted against the temperature. The red circles and lines show individual cells, the black circles and line the mean and S.E.M. of the 5 cells.

To test the possible involvement of intracellular factors in the temperature activation mechanism, we measured TREK-1 channels in different patch configurations. Importantly, the observed thermosensitivity was preserved upon prolonged heating in on-cell measurements, where the current maximum was stable at constant temperature (Fig. 2B; fold change 8.50 ± 1.17). In contrast, after excision of the patches, yielding the inside-out configuration, channels immediately and completely lost their temperature sensitivity (Fig. 2B). When measurements were done in the inside-out configuration from the beginning, the channels display no thermosensitivity (Fig. 2C; fold change 1.2 ± 0.25), but still were functional as they were activated by other stimuli, i.e. arachidonic acid or BL-1249 (Fig. 2C). Hence, together with the delay in the onset of current activation after temperature rise, this dependence on cell integrity suggested that temperature activation is not an intrinsic channel property.

Since we so far used a broad temperature range between 10-50 °C to characterize temperature activation of the current, the question arose whether TREK channels are activated in the physiologically relevant temperature range. Therefore, we measured TREK-1 channels in the on-cell configuration at 35 °C and further raised the temperature to about 40 °C. Indeed, a small but clear temperature-dependent current increase was induced (Fig. 2E). Plotting the current increase induced by a temperature rise from room temperature to 35 °C and the rise from 35 °C to 40 °C yielded a linear relationship, meaning that the temperature sensor of TREK-1 channels was equally active between 21-35 °C and at temperatures around normal body temperature, i.e. 35-40 °C (Fig. 2F).

### The C-terminal domain of TREK-1 channels contains a temperature-responsive element

We intended to identify the channel region responsible for the thermosensitivity in TREK channels. It has been reported previously that the C-terminus of TREK-1 channels is indispensable for their temperature activation, and a decoupling between the CTD and the pore-forming core also leads to the loss of thermosensitivity (Maingret *et al*., 2000; Bagriantsev, Clark and Minor, 2012). We have shown above that TRAAK channels were not thermosensitive (Fig. 1 and S2). In order to prove the importance of the TREK C-terminus for temperature activation, we constructed chimeric channels where the entire C-termini were swapped between the two channels (after residues V313 of TREK-1 and V259 of TRAAK), yielding the constructs TREK-1/ctTRAAK and TRAAK/ctTREK-1, respectively (Fig. 3A). Indeed, the swapping of the C-termini led to an exchange of their thermosensitivity. While TREK-1/ctTRAAK channels entirely lost their thermosensitivity with the introduction of the TRAAK C-terminus (fold change 2.53 ± 0.60), thermosensitivity was transferred to TRAAK/ctTREK-1 channels with the C-terminus of TREK-1, with an activation comparable to the TREK-1 wildtype channel (Fig. 3A, B; fold change 12.45 ± 2.65 for WT TREK-1 and 10.77 ± 2.01 for TRAAK/ctTREK-1). This transfer of the thermosensitivity by connecting the respective pore forming core to the TREK-1 C-terminus was only effective within the TREK/TRAAK subfamily, as other chimeric channels failed to yield temperature sensitive channels (i.e. TALK-2 or TASK-3 core channels with TREK-1 C-terminus; fold change of 2.81 ± 0.75 for TALK-2/ctTREK-1 channels and 2.13 ± 0.46 for TASK-3/ctTREK-1 channels; Fig. 3B).

**Figure 3.**
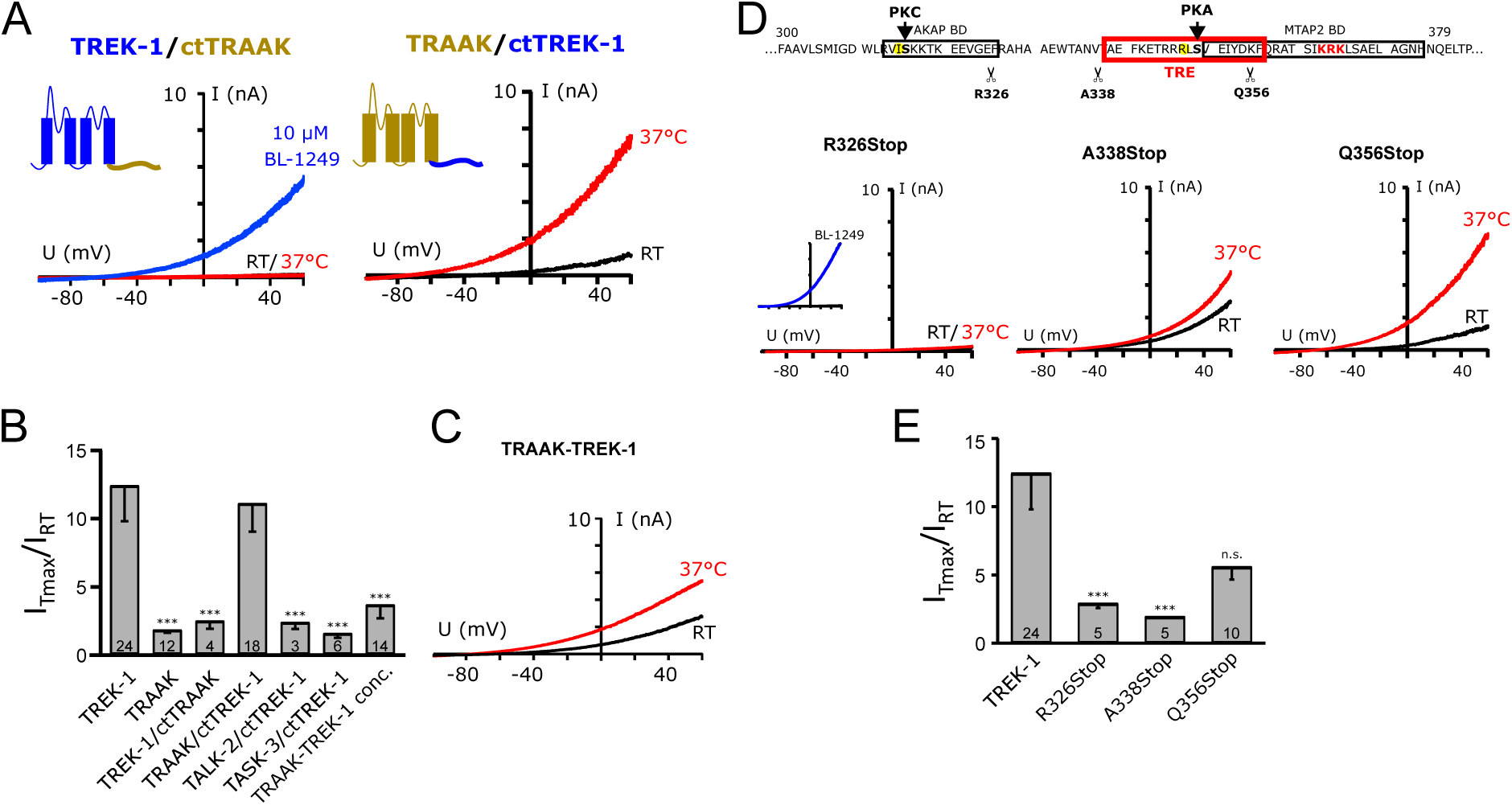
Identification of the temperature responsive element (TRE) in the C-terminus of TREK-1 channels. **A** Exemplary current traces of HEK293 cells transfected with chimeric constructs of TREK-1 core channels with C-termini of TRAAK channels (TREK-1/ctTRAAK, left) and vice versa (TRAAK/ctTREK-1, right) measured at RT (black trace) and 37 °C (red trace). 10 µM BL-1249 (blue trace) was used to show the functionality of the chimeric TREK-1/ctTRAAK channels. **B** The bar diagram shows the fold change of channel activation upon temperature elevation for TREK-1 and TRAAK wild type and different chimeric channels. **C** Exemplary current traces of concatenated TRAAK-TREK-1 channels measured at RT (black trace) and 37 °C (red trace). **D** Part of the TREK-1 C-terminal sequence with the known PKC and PKA phosphorylation sites. The binding motifs for AKAP150 and MAP2 are marked with black boxes, the identified TRE with a red box. The scissors indicate the sites of truncation. Below, exemplary current traces of the truncated TREK-1 channels are shown (black traces: at RT; red traces: at 37 °C; inset: control with 10 µM BL-1249). **E** Bar diagram showing the fold change of activation upon temperature elevation for truncated TREK-1 channels.

As K2P channels form dimers to build functional channels, homomeric channels have two identical C-termini. To investigate the influence of one thermosensitive C-terminus on the thermosensitivity of the whole channel, we constructed concatenated channels of TREK-1 subunits with temperature-insensitive TRAAK channel subunits. Interestingly, concatemers of TRAAK and TREK-1 subunits were not activated by temperature, providing an indication that one TREK-1 subunit is not enough for temperature-sensitivity (fold change 3.40 ± 0.81, Fig. 3B, C).

To further narrow down the region responsible for thermosensitivity, we gradually truncated the C-terminus of TREK-1 (Fig. 3D). All truncated constructs were still functional and regulated by BL-1249 (inset for R326Stop in Fig. 3D). Truncating the complete TREK-1 C-terminus at arginine 326 (R326Stop) led to a complete loss of thermosensitivity (fold change of 2.9 ± 0.33, Fig. 3D, E). The thermosensitivity was still lost upon truncation at alanine 338 (A338Stop, fold change 1.93 ± 0.07). In contrast, the truncation of only the last 70 amino acids (Q356Stop) had little effect on the thermosensitivity of TREK-1 channels (fold change 5.64 ± 0.91). Thus, the stretch of 18 amino acids from A338 to F355 must comprise the temperature responsive element of TREK-1 channels (TRE hereinafter).

### The PKA-mediated phosphorylation state regulates thermosensitivity

In addition to the known phosphorylation sites for PKC and PKA, the TREK-1 C-terminus contains binding sites for the interacting proteins AKAP5 (AKAP150 in mouse) and microtubule associated protein 2 (MAP2) close-by (Fig. 3D; Sandoz et al. 2006 and 2008). A sequence analysis of the TRE revealed that two of these previously described modifiable elements lie within this region: the PKA phosphorylation site at S348 and the MAP2 binding domain from E350-Q375. TREK-1 channels are modulated by phosphorylation by PKC and PKA at two serines (S315 and S348; (Patel *et al*., 1998; Murbartián *et al*., 2005). Therefore, we sought to investigate the influence of the kinases on the thermosensitivity of TREK-1 channels using pharmacological modulators. The PKA site S348 (S333 in the shorter isoform) in TREK-1 is also present in the other thermosensitive K2P channel TREK-2 (here S364), while it is absent in the thermoinsensitive TRAAK channels (Fig. S2A). In contrast, the PKC phosphorylation site S315 (S300 in the shorter isoform) is present in TREK-2 (S331) as well as TRAAK (S287), though it has been described to be not functional in TRAAK channels (Fink *et al*., 1998).

Activation of PKC by phorbol-12-myristate-13-acetate (PMA; 1 µM), though reducing the basal current, had little effect on temperature activation (fold change 13.50 ± 3.25; Fig. 4). In contrast, the activation of PKA by 10 µM forskolin/100 µM 3-isobutyl-1-methylxanthine (Forsk/IBMX) dramatically inhibited the basal current and additionally abolished the temperature activation of TREK-1 channels (fold change 3.21 ± 0.54; Fig. 4). This was not due to dysfunction of the channels as they were still activated pharmacologically by BL-1249. Preincubation with the PKA inhibitor H-89 completely prevented this inhibitory effect of Forsk/IBMX on the thermosensitivity (fold change 12.89 ± 6.07; Fig. 4 insets). Inhibition of PKA by 1 µM H-89 alone had no effect or even augmented the temperature activation (fold change of 18.00 ± 6.99; Fig. 4D). Hence, thermosensitivity of TREK-1 channels is specifically mediated by PKA phosphorylation, and PKA activation is sufficient to disable a temperature response. Importantly, this PKA mediated phosphorylation itself could not constitute the temperature dependent step, as the thermosensitivity was preserved after inhibition of PKA by H89.

**Figure 4.**
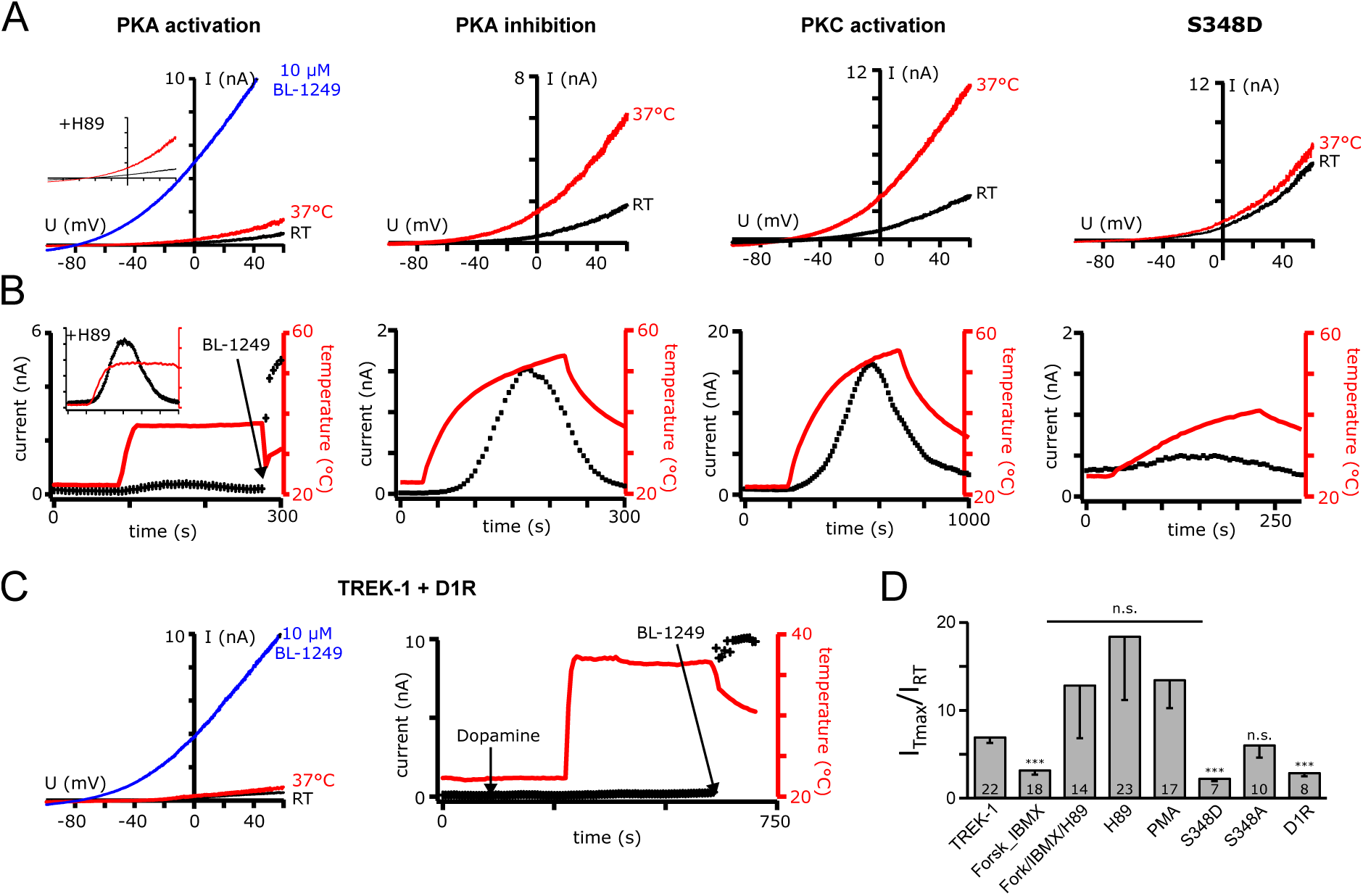
PKA phosphorylation specifically suppresses thermosensitivity in TREK-1 channels. **A** Exemplary basal and temperature activated TREK-1 currents after pharmacological kinase modulation. The inset shows an example of a cell preincubated with H-89 before the application of Forsk/IBMX (black traces: at RT; red traces: at 37 °C). **B** Time courses of temperature activated currents with kinase modulation as shown in A. **C** Exemplary current traces and time course of TREK-1 currents co-expressed with the G protein coupled dopamine receptor 1 (Drd1) exposed to dopamine and temperature elevation. As no currents were activated, the channels were activated by 10 µM BL-1249 after temperature elevation (blue trace/ arrow). **D** Bar diagram showing the fold change of temperature activation after kinase modulation or Drd1 receptor activation.

In addition, we mutated the PKA phosphorylation site in TREK-1 channels either to aspartate to mimic the phosphorylated state (S348D) or to alanine to mimic the dephosphorylated state (S348A). Like the activation of PKA by Forsk/IBMX, the S348D mutation led to a dramatic decrease in basal current and completely abolished the temperature activation, supporting the importance of the degree of PKA-mediated phosphorylation for the thermosensitivity of TREK channels (fold change: 2.27 ± 0.34; Fig. 4). Likewise, in agreement with the inhibition of PKA by H-89, the S348A mutation had no effect on the thermosensitivity (fold change 6.07 ± 1.46; Fig. 4D).

Finally, we activated PKA via a physiological pathway by coexpression of dopamine receptor 1 (Drd1) as an example of a G_αs_ coupled receptor. Consistently, PKA activation via the receptor mediated cAMP synthesis prohibited temperature activation and reduced temperature activated currents compared to cells not expressing Drd1 (fold change 2.92 ± 0.43, Fig 4C, D).

### Contact to the cytoskeleton is necessary for temperature activation

An interaction between MAP2 and the MAP2 binding site in TREK-1 (E350-Q375) has been shown by co-immunoprecipitation experiments with synaptosomal brain proteins(Sandoz *et al*., 2008). As MAP2 links the associated interaction partners to the cytoskeleton, the overlap of the MAP2 binding site with the TRE (E339-Q356) suggested a possible involvement of MAP2 mediated contact between TREK-1 channels and the cytoskeleton in temperature sensing. To identify which of the different parts of the cytoskeleton is linked to the TREK-1 C-terminus, we used colchicine (0.5 mM) and vinblastine (50 µM) to disrupt the microtubule network, and cytochalasin B (25 µM) to disrupt the polymerization of the actin cytoskeleton (Fig. 5A, B).

**Figure 5.**
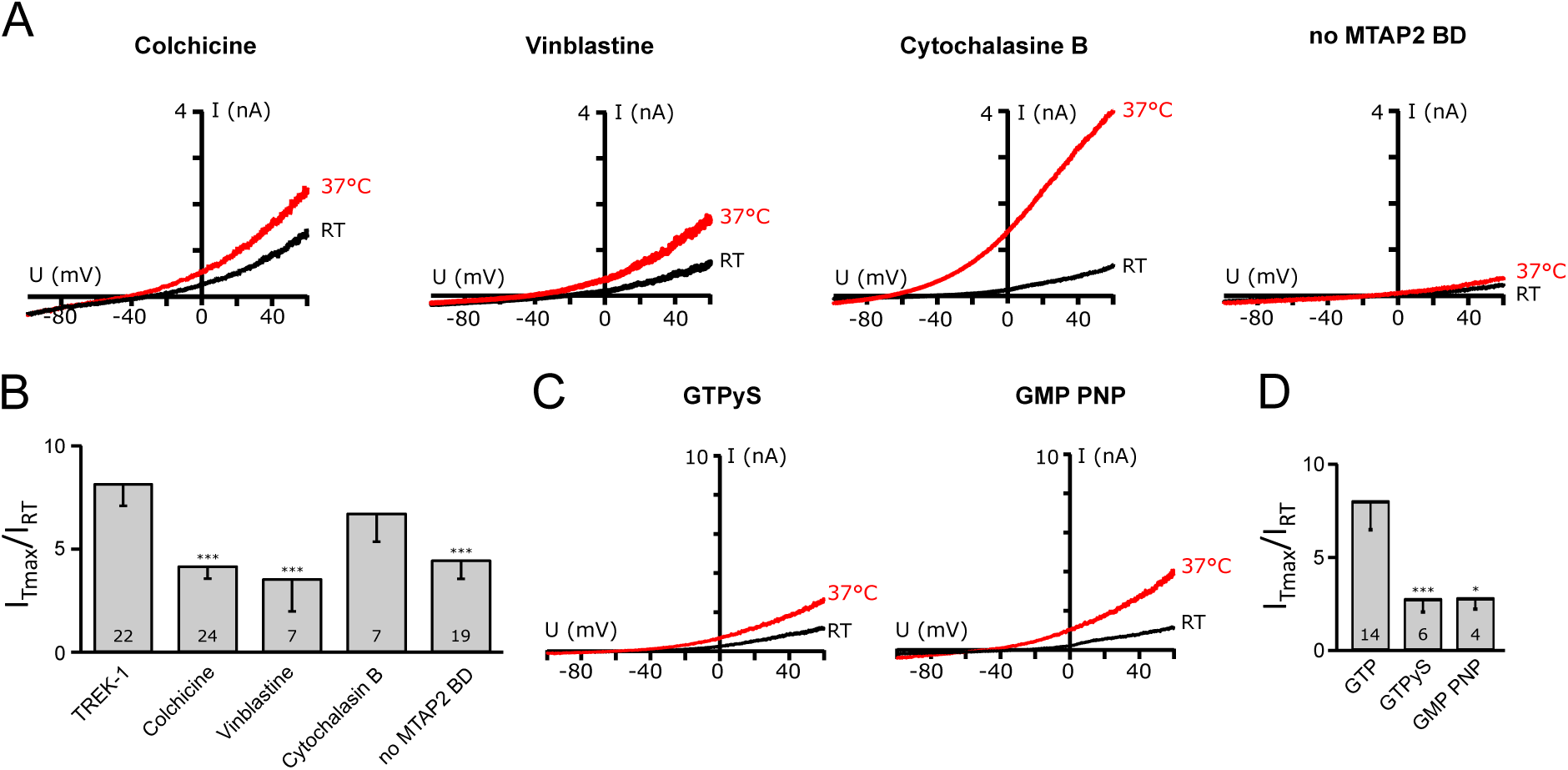
Thermosensitivity of TREK-1 channels depends on the connection of the C-terminus to the microtubular network. **A** Exemplary temperature stimulated TREK-1 currents after pharmacological modulation of the cytoskeleton or mutation of the MAP2 binding motif (black traces: at RT; red traces: at 37 °C). **B** The bar diagram shows the fold change of temperature activation from measurements as in A analyzed at +60 mV. **C** Current traces of TREK-1 channels measured with the non-hydrolysable GTP analogs GTPγS or GMP PNP in the pipette solution (black traces: at RT; red traces: at 37 °C). **D** Bar diagram showing the fold change of temperature activated currents in the presence of GTP and its non-hydrolysable analogs.

Interestingly, interference with the actin cytoskeleton had only a minor effect on temperature activation (fold change 6.74 ± 1.39), while the disassembly of the microtubular network led to the suppression of the temperature activation (fold changes 4.18 ± 0.61 and 3.56 ± 1.58 for colchicine and vinblastine, respectively).

As for the formation of microtubule the hydrolysis of GTP bound to tubulin is necessary, we also included non-hydrolysable GTP analogs in our pipette solution. Neither in the presence of GTPγS nor with GMP PNP instead of GTP we measured any temperature activated TREK-1 current (Fig. 5C, D; fold change for GTP 8.02 ± 1.53, for GTPγS 2.76 ± 0.7, and for GMP PNP 2.74 ± 0.68).

Sandoz et al. (2008) have shown that an exchange of the amino acids R357, K362, R363, and K364 to proline impedes the TREK-1/MAP2 interaction. To test for the contribution of the TREK-1/MAP2 interaction to the temperature sensitivity, we mutated the MAP2 binding site. This significantly reduced the thermosensitivity of the channels (Fig. 5A, B; fold change 4.47 ± 0.92), again suggesting that the integrity of the microtubules is necessary for temperature sensitivity of TREK-1 channels.

## Discussion

In this study, we report the first localized TRE in a mammalian thermosensitive ion channel. We identified a region in the C-terminus of the channel as being indispensable for its thermosensitivity. Especially, the PKA phosphorylation site and the MAP2 binding site are essential for temperature-activation of TREK channels.

In mammals, temperature perception is mainly mediated by temperature-sensitive ion channels, primarily by members of the TRP channel family. Despite intensive efforts, the molecular basis for their thermosensitivity has not yet been fully elucidated. Two general mechanisms of thermosensitivity are discussed: global vs. localized mechanisms (Arrigoni and Minor, 2018). A hypothesis that represents a global mechanism is the temperature-dependent change in specific heat capacity of a channel (Clapham and Miller, 2011; Yeh, Jara-Oseguera and Aldrich, 2023). This would enable temperature-dependent gating transitions without the need of specific TREs, and the overall heat capacity of open and closed states could also be altered by e.g. binding events, mutations or deletions in distant parts of the channel. For example, Chowdhury, Jarecki and Chanda (2014) were able to obtain a temperature-gated Shaker K_v_ channel by rational design of amino acid positions involved in gating transitions, emphasizing the importance of solvation changes for the heat capacity of the channel. In contrast, a localized mechanism would require the existence of a defined TRE that controls the channel gate in response to its stimulus. Examples of both can be found in temperature-sensitive ion channels. In most thermoTRP channels, changes in very different parts of the protein impede thermosensitivity, and no distinct temperature-sensing element could be identified to date. In the well-studied TRPV1 channel, the pore region seems to be essential as mutations or deletions in the extracellular part of S6 and the pore (Grandl *et al*., 2008, 2010), or the pore turret (Yang *et al*., 2010; Cui *et al*., 2012) impair its temperature-sensitivity. Furthermore, Zhang *et al*. (2018) were able to transfer the temperature-sensitivity from TRPV1 to the Shaker K_v_ channel simply by exchanging their pore domains, supporting the idea of a localized thermosensor in these channels. Contradictory to this, intracellular parts of the TRP channels, e.g. the membrane proximal domain between the ankyrin repeat domain (ARD) and TM1 (Yao, Liu and Qin, 2011) and the C-terminus (Brauchi *et al*., 2006) have also been found to influence its temperature sensitivity in TRP channels. In bidirectional heat- and cold-activated TRPA1 channel, deletions in the N-terminal voltage sensing like domain (VSLD) abolished exclusively heat sensitivity, but cool sensitivity seemed to be distributed to linker and CTD with possible contributions of N-terminal ARD and the VSLD domains (Jabba *et al*., 2014; Moparthi *et al*., 2022). In addition, recent cryo-EM structures of TRPV1 and TRPV3 channels generated at different temperatures also point to a global temperature activation in these channels (Singh *et al*., 2019; Kwon *et al*., 2021; Nadezhdin *et al*., 2021). To our knowledge, a clearly delimited area of the protein which represents a true localized mechanism of temperature sensitivity could only be identified in one ion channel: In the bacterial Na_v_ channel of *Silicibacter pomeroyi*, a short ≈14 amino acid stretch in a metastable domain connecting the CTD to the pore was shown to control temperature-dependent voltage gating depending on its degree of disorder (Arrigoni *et al*., 2016).

Several studies have shown that TREK channels are temperature sensitive, but the mechanism of temperature activation remained unknown (Maingret *et al*., 2000; Kang, Choe and Kim, 2005; Noël *et al*., 2009; Bagriantsev, Clark and Minor, 2012; Pereira *et al*., 2014; Viatchenko-Karpinski, Ling and Gu, 2018). In agreement with previous investigations, we measured Q10 values up to 20 for TREK-1 and TREK-2 channels (Maingret *et al*., 2000; Kang, Choe and Kim, 2005). All other tested representatives from the different K2P subfamilies were not temperature activated and exhibited Q10 values below 4. Surprisingly, the third member of this K2P subfamily, TRAAK, which is closely related to TREK-1 and TREK-2, was not activated by temperature. It has been postulated that TRAAK channels are thermosensitive, albeit with a higher activation threshold (31 °C) than TREK-1 and TREK-2 channels (Kang, Choe and Kim, 2005). However, we did not find any thermosensitivity upon heating to temperatures of 40 °C and more though we tested human and murine TRAAK channels in both expression systems (HEK293 and COS7 cells).

The C-terminus of TREK-1 channels is of great importance for the integration of various stimuli in addition to thermosensitivity (Maingret *et al*., 2000), including intracellular pH (Maingret *et al*., 1999), polyunsaturated fatty acids (Patel *et al*., 1998), PIP_2_ (Chemin *et al*., 2005), and phosphorylation (Fink *et al*., 1996; Murbartián *et al*., 2005), and it provides binding sites for interacting proteins (Sandoz *et al*., 2006, 2008). By swapping the TRE-containing C-terminus of TREK-1 with that of the non-thermosensitive TRAAK channel we were able to transfer the thermosensitivity to the TRAAK channel, and vice versa make TREK-1 channels temperature-insensitive. This shows that the TREK-1 C-terminus is not only necessary but sufficient for temperature sensitivity within the TREK/TRAAK subfamily. Interestingly, the heteromeric channel formed of one TRAAK subunit and one TREK-1 subunit was not temperature-sensitive, suggesting that two TREK C-termini are needed for temperature activation. In good agreement, Bagriantsev, Clark and Minor (2012) have shown that a concatemer of two TREK-1 subunits where one had an uncoupled C-terminus had a blunted temperature response. By further step-wise truncation, we consequently isolated the TRE critical for temperature activation in the TREK-1 CTD to 18 amino acids (A338-F355). Importantly, this region includes the PKA phosphorylation site and overlaps with the MAP2 binding site which connects the TREK-1 C-terminus to the cytoskeleton.

In general, temperature-sensitive channels differ in whether they are intrinsically temperature-sensitive or require a co-factor. For example, TRPV1, TRPV3, or TRPM8 channels are still regularly temperature-sensitive after reconstitution in lipid bilayers, so it can be assumed that these channels are intrinsically temperature-dependent (Zakharian, Cao and Rohacs, 2010; Cao *et al*., 2013; Nadezhdin *et al*., 2021). In contrast, TRPV4 channels, like TREK-1 channels, lose their temperature-sensitivity when their cellular context is removed (Watanabe *et al*., 2002). The fact that temperature activation is absent when TREK channels are measured in excised patches (Maingret *et al*., 2000; Kang, Choe and Kim, 2005), and our observation that temperature activated currents decrease in whole-cell measurements and upon inside-out patch excision is consistent with the loss of intracellular factors that convey thermosensitivity. Furthermore, the time delay of activation after the inset of temperature rise seen in TREK-1 may hint at the involvement of additional processes, while TRPV1 activity closely follows the rate of temperature elevation (Cao *et al*., 2013). To shed light on the mechanism of TREK-1 temperature activation, we investigated the role of different components of the cytoskeleton as putative MAP2 binding partners as well as the relevance of kinase phosphorylation. Appropriately, we found that microtubule dynamics determined the temperature activation of TREK-1 channels. When we pharmacologically manipulated different cytoskeleton constituents, only the disruption of the microtubules or interference with their synthesis, i.e. by non-hydrolysable GTP or GTP-PNP, impeded thermosensitivity.

Temperature has a strong influence on the equilibrium between catastrophe and rescue of microtubule (how disassembly and assembly are called), with higher temperatures promoting the rescue. Significantly, MAP proteins regulate dynamics of microtubules in a process called selective stabilization and protect e.g. from cold-induced disassembly (reviewed by Dehmelt and Halpain, 2004; Baas *et al*., 2016; DeGiosio *et al*., 2022). The family member MAP2 is expressed in neurons and was shown to co-immunoprecipitate with TREK-1 channels in a lysate obtained from mouse brains (Sandoz *et al*., 2008; Andharia *et al*., 2017). Our results show that the disconnection of MAP2 and TREK-1 channels by mutating the MAP2 binding region in the C-terminus of TREK-1 channels completely suppressed the thermosensitivity. This means that a change in temperature leads to a modified connection between the TRE of TREK-1 and the microtubular network, which then causes the activation of TREK-1 channels with increasing temperatures, and their closure upon cooling.

Interestingly, beyond modulation of the fundamental temperature dependence of assembly/ disassembly, the family member MAP6 has been described as temperature sensor that binds to microtubules at lower temperatures (Delphin *et al*., 2012). Mechanistically, a linear unfolding of a specific domain from beta structures at high temperatures to a disordered state at low temperatures was shown, a structural feature which is also present in the MAP2/tau subfamily (Delphin *et al*., 2012). Furthermore, this non-intrinsic temperature activation of TREK-1 channels is modulated by the current state of the cell, reflected by PKA activity. The pharmacological or receptor-mediated activation of PKA specifically abolished temperature activation, while PKC activation, though also inhibiting the basal current, had no effect on the temperature dependence of the channel. The prevention of temperature activation by PKA-mediated phosphorylation has been postulated previously by Maingret *et al*. (2000), who used cAMP to inhibit the channels at different temperatures. The inhibition was absent in the PKA site mutant S333A (S348A in the TREK-1 isoform used in this study), and was interpreted as a reversal of the temperature activation by the activation of PKA. However, the PKA-mediated phosphorylation itself cannot be the temperature sensitive step, as otherwise the inhibition of PKA by H-89 or the dephosphorylation mimicking mutant S348A should also have led to the suppression of thermosensitivity in our experiments. Thus, the phosphorylation of TREK-1 channels by PKA only changes the status of the channel, rendering it insensitive to temperature activation. Phosphorylation operates as a molecular switch, which allows the cell to adapt the TREK-1 temperature sensitivity by changing the degree of PKA-mediated phosphorylation, thereby coupling it to active signaling pathways.

The contact between the C-terminus of TREK-1 channels, the cytoskeleton, and the catalytic domain of PKA is likely mediated by MAP2 (Obar *et al*., 1989; Melková *et al*., 2019). Therefore, MAP2 is a key player in the temperature-dependent activation of TREK-1 channels and critical in the recruitment of PKA to the TREK-1 C-terminus. When S348 is dephosphorylated, the channel is connected to the microtubular network and temperature-sensitive; upon phosphorylation, the connection may be altered or lost, and the channel is no longer sensitive to temperature.

It has been shown that in TREK channels, mechanoactivation induces a conformational change from a low activity ‘down’ conformation to a high activity ‘up’ conformation, where the proximal CTD is close to the inner membrane leaflet (Brohawn, Su and MacKinnon, 2014; Dong *et al*., 2015). This has been postulated also for other stimuli including temperature and dephosphorylation (Honoré, 2007; McClenaghan *et al*., 2016; Türkaydin *et al*., 2024). One may speculate that the MAP2-mediated contact to the cytoskeleton stabilizes TREK-1 channels in the temperature-induced high activity ‘up’ conformation (McClenaghan *et al*., 2016). Phosphorylation, however, would favor the low-activity ‘down’ state (Türkaydin *et al*., 2024), switching off temperature activation.

TREK channels are expressed in both DRG/TG neurons and thermosensitive neurons of the POA (Maingret *et al*., 2000; Talley *et al*., 2001; Yamamoto, Hatakeyama and Taniguchi, 2009; Marsh *et al*., 2012; Viatchenko-Karpinski, Ling and Gu, 2018) and may therefore contribute to peripheral and central thermosensation. In DRG/TG neurons, TREK-1 is the counterpart of TRPV1 channels, and accordingly, these neurons would be medium temperature sensitive at rest, as their stimulation by TRPV1 activation would be dampened by TREK-1 channel activation. However, TRPV1 channels are sensitized to temperature changes by PKA-mediated phosphorylation, e.g. in ongoing inflammation (Bhave *et al*., 2002; Fischer and Reeh, 2007), where mediators like prostaglandins activate PKA (Hucho and Levine, 2007), and, correspondingly, many studies have connected heat hyperalgesia during inflammation to TRPV1 channels (Vriens, Nilius and Voets, 2014). Conversely, TREK-1 channels are desensitized to temperature changes by PKA-mediated phosphorylation (Maingret *et al*., 2000 and present study), and the suppression of TREK channel thermosensitivity may as well contribute to prostaglandin mediated hyperalgesia. Thus, the firing threshold of thermoreceptor fibers might be controlled by a balance of the phosphorylation states of both antagonistic channels.

In the POA the core body temperature is closely monitored, but the set point for body temperature is not fixed, e.g. during sleep the set point is lowered, whereas it is elevated in fever (Tan and Knight, 2018; Nakamura, 2024). Fever induced by pathogens like bacteria is mediated by pyrogens as prostaglandin E2, which suppresses the cAMP/PKA pathway through G_αi_ coupled EP_3_ prostaglandin receptors. It has been established that EP_3_ receptor activation results in the decrease of the firing rate in the subset of ‘warm-sensitive’ POA neurons, leading to subsequent thermogenesis (Ranels and Griffin, 2003, 2005; Morrison, Nakamura and Madden, 2008). Model simulations of such neurons showed that the expression of temperature-activated TREK channels is capable of reducing their firing rate in the physiological temperature range (32-40 °C; Wechselberger *et al*., 2006). Therefore, TREK thermoactivation upon suppression of PKA-mediated phosphorylation could contribute to the change in temperature setpoint of POA neurons under inflammatory conditions.

---

**Table S1.**
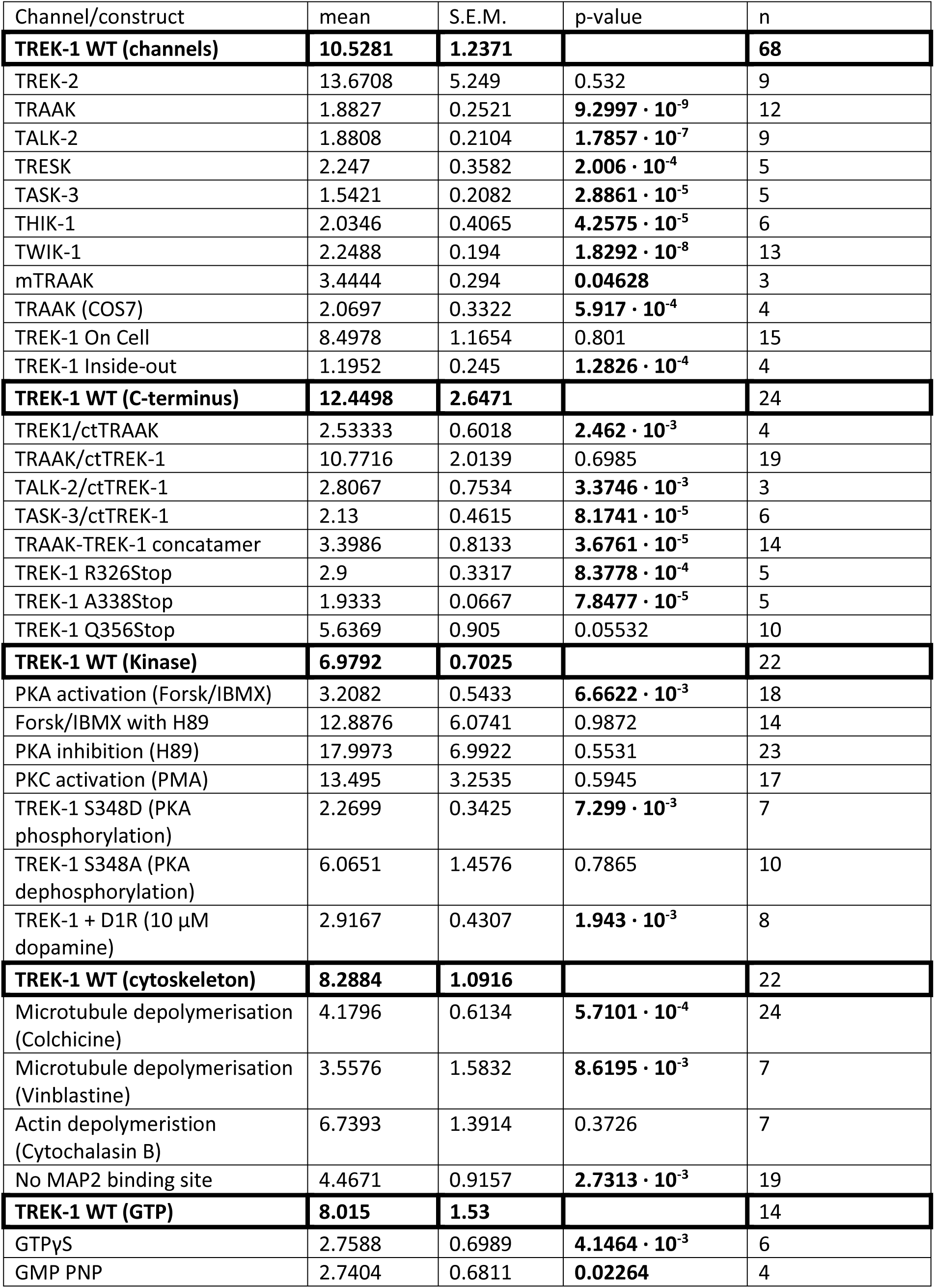
Maximal current increase by temperature elevation. . Fold change of currents relative to basal current at RT were determined with the ramp protocol at +60 mV. Different control values for TREK-1 WT are given for the different data sets. P-values highlighted in bold are significant.

| Channel/construct | mean | S.E.M. | p-value | n |
| --- | --- | --- | --- | --- |
| **TREK-1 WT (channels)** | **10.5281** | **1.2371** |  | **68** |
| TREK-2 | 13.6708 | 5.249 | 0.532 | 9 |
| TRAAK | 1.8827 | 0.2521 | **9.2997 · 10<sup>-9</sup>** | 12 |
| TALK-2 | 1.8808 | 0.2104 | **1.7857 · 10<sup>-7</sup>** | 9 |
| TRESK | 2.247 | 0.3582 | **2.006 · 10<sup>-4</sup>** | 5 |
| TASK-3 | 1.5421 | 0.2082 | **2.8861 · 10<sup>-5</sup>** | 5 |
| THIK-1 | 2.0346 | 0.4065 | **4.2575 · 10<sup>-5</sup>** | 6 |
| TWIK-1 | 2.2488 | 0.194 | **1.8292 · 10<sup>-8</sup>** | 13 |
| mTRAAK | 3.4444 | 0.294 | **0.04628** | 3 |
| TRAAK (COS7) | 2.0697 | 0.3322 | **5.917 · 10<sup>-4</sup>** | 4 |
| TREK-1 On Cell | 8.4978 | 1.1654 | 0.801 | 15 |
| TREK-1 Inside-out | 1.1952 | 0.245 | **1.2826 · 10<sup>-4</sup>** | 4 |
| **TREK-1 WT (C-terminus)** | **12.4498** | **2.6471** |  | **24** |
| TREK1/ctTRAAK | 2.53333 | 0.6018 | **2.462 · 10<sup>-3</sup>** | 4 |
| TRAAK/ctTREK-1 | 10.7716 | 2.0139 | 0.6985 | 19 |
| TALK-2/ctTREK-1 | 2.8067 | 0.7534 | **3.3746 · 10<sup>-3</sup>** | 3 |
| TASK-3/ctTREK-1 | 2.13 | 0.4615 | **8.1741 · 10<sup>-5</sup>** | 6 |
| TRAAK-TREK-1 concatamer | 3.3986 | 0.8133 | **3.6761 · 10<sup>-5</sup>** | 14 |
| TREK-1 R326Stop | 2.9 | 0.3317 | **8.3778 · 10<sup>-4</sup>** | 5 |
| TREK-1 A338Stop | 1.9333 | 0.0667 | **7.8477 · 10<sup>-5</sup>** | 5 |
| TREK-1 Q356Stop | 5.6369 | 0.905 | 0.05532 | 10 |
| **TREK-1 WT (Kinase)** | **6.9792** | **0.7025** |  | **22** |
| PKA activation (Forsk/IBMX) | 3.2082 | 0.5433 | **6.6622 · 10<sup>-3</sup>** | 18 |
| Forsk/IBMX with H89 | 12.8876 | 6.0741 | 0.9872 | 14 |
| PKA inhibition (H89) | 17.9973 | 6.9922 | 0.5531 | 23 |
| PKC activation (PMA) | 13.495 | 3.2535 | 0.5945 | 17 |
| TREK-1 S348D (PKA phosphorylation) | 2.2699 | 0.3425 | **7.299 · 10<sup>-3</sup>** | 7 |
| TREK-1 S348A (PKA dephosphorylation) | 6.0651 | 1.4576 | 0.7865 | 10 |
| TREK-1 + D1R (10 µM dopamine) | 2.9167 | 0.4307 | **1.943 · 10<sup>-3</sup>** | 8 |
| **TREK-1 WT (cytoskeleton)** | **8.2884** | **1.0916** |  | **22** |
| Microtubule depolymerisation (Colchicine) | 4.1796 | 0.6134 | **5.7101 · 10<sup>-4</sup>** | 24 |
| Microtubule depolymerisation (Vinblastine) | 3.5576 | 1.5832 | **8.6195 · 10<sup>-3</sup>** | 7 |
| Actin depolymerisation (Cytochalasin B) | 6.7393 | 1.3914 | 0.3726 | 7 |
| No MAP2 binding site | 4.4671 | 0.9157 | **2.7313 · 10<sup>-3</sup>** | 19 |
| **TREK-1 WT (GTP)** | **8.015** | **1.53** |  | **14** |
| GTPγS | 2.7588 | 0.6989 | **4.1464 · 10<sup>-3</sup>** | 6 |
| GMP PNP | 2.7404 | 0.6811 | **0.02264** | 4 |

**Figure S1.**
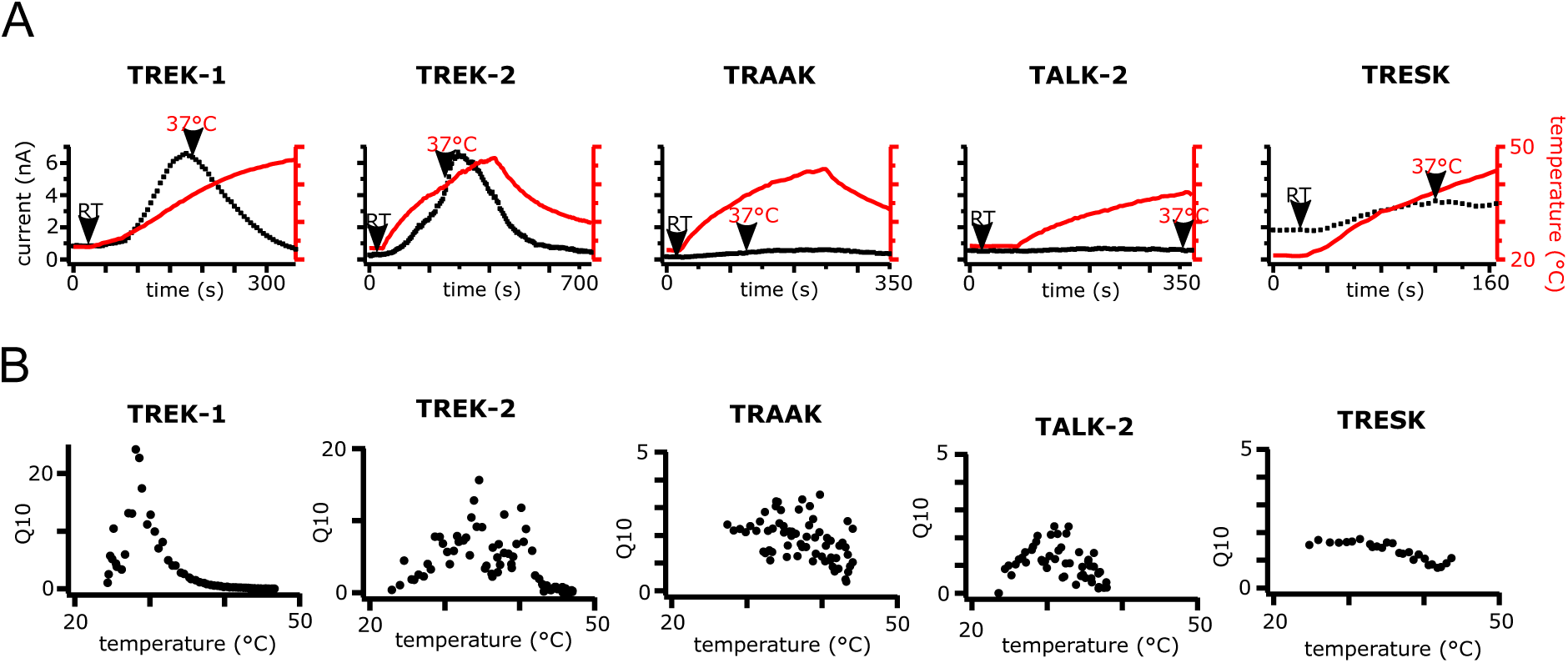
The Q10 values of TREK channels are temperature-dependent. **A** Time courses of the current response of different tested K2P channels to temperature elevation. The currents were elicited with a ramp protocol and plotted at 0 mV. The red line shows the time course of the monitored bath temperature; the arrow indicates the current at 37 °C. **B** Exemplary Q10 values for the currents shown in A. The values were calculated for each pair of ramps during the temperature elevation, yielding Q10 values up to 20 for TREK-1 and TREK-2. For the other tested K2P channels maximal Q10 values of 3 were calculated (note the different scaling on the y axes).

**Figure S2.**
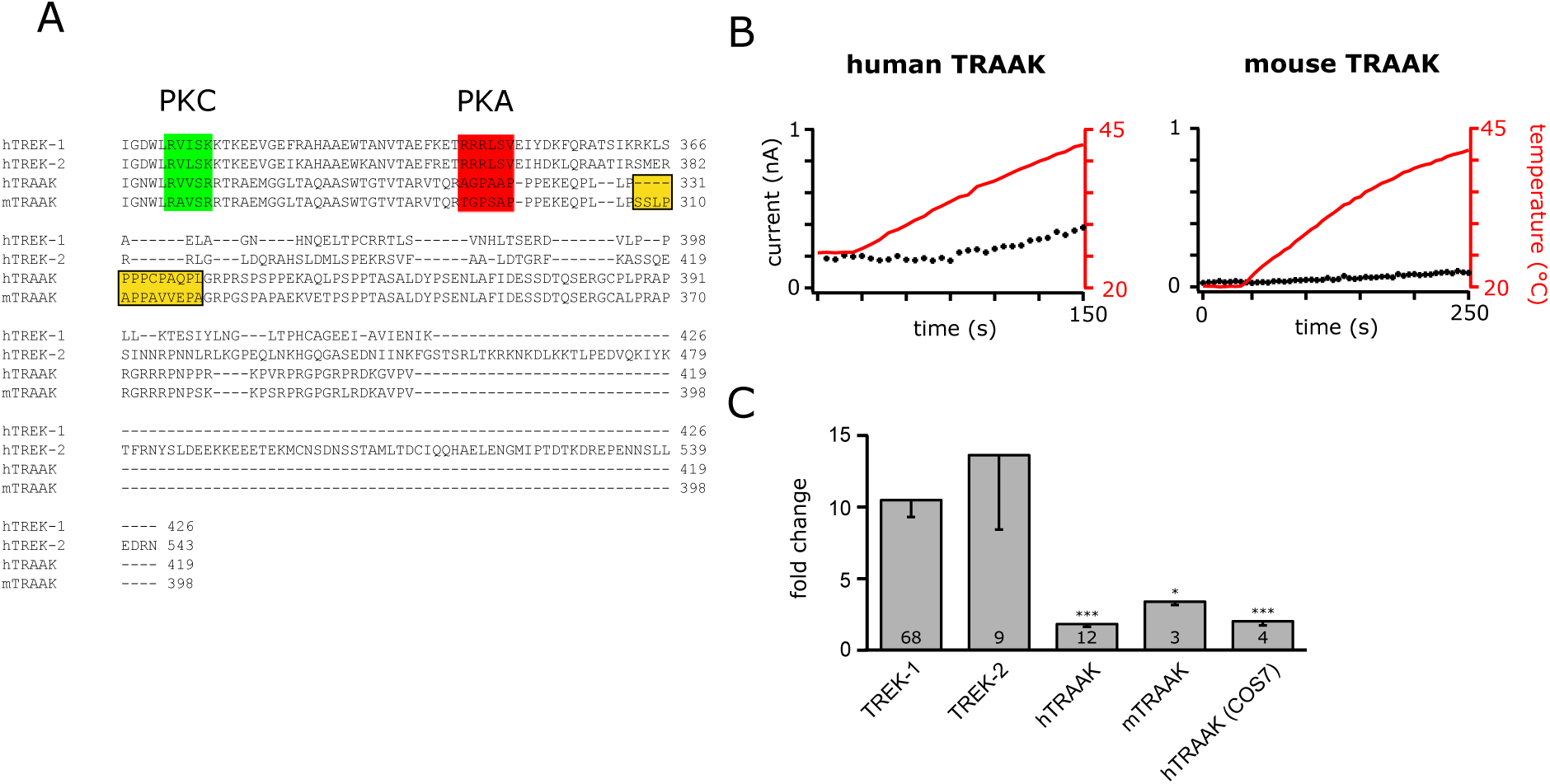
Neither human nor murine TRAAK channels are thermosensitive. **A** Sequence alignment of the C termini of human K2P members of the TREK/TRAAK subfamily and mouse TRAAK. Highlighted are the phosphorylation sites for PKC (green, conserved in TRAAK) and PKA (red, not conserved in TRAAK). In addition, a stretch of 9 amino acids in human TRAAK (13 amino acids in mouse) is highlighted that is not conserved between human and murine TRAAK (orange). **B** Time course of current changes upon temperature elevation in human and mouse TRAAK channels. Currents were measured with a ramp protocol and the currents at 0 mV were plotted. Red lines: changes in bath temperature measured in °C. **C** Fold change of TREK-1 and TREK-2 currents induced by temperature elevation compared to human and mouse TRAAK channels expressed in HEK293 and COS7 cells.

**Figure S3.**
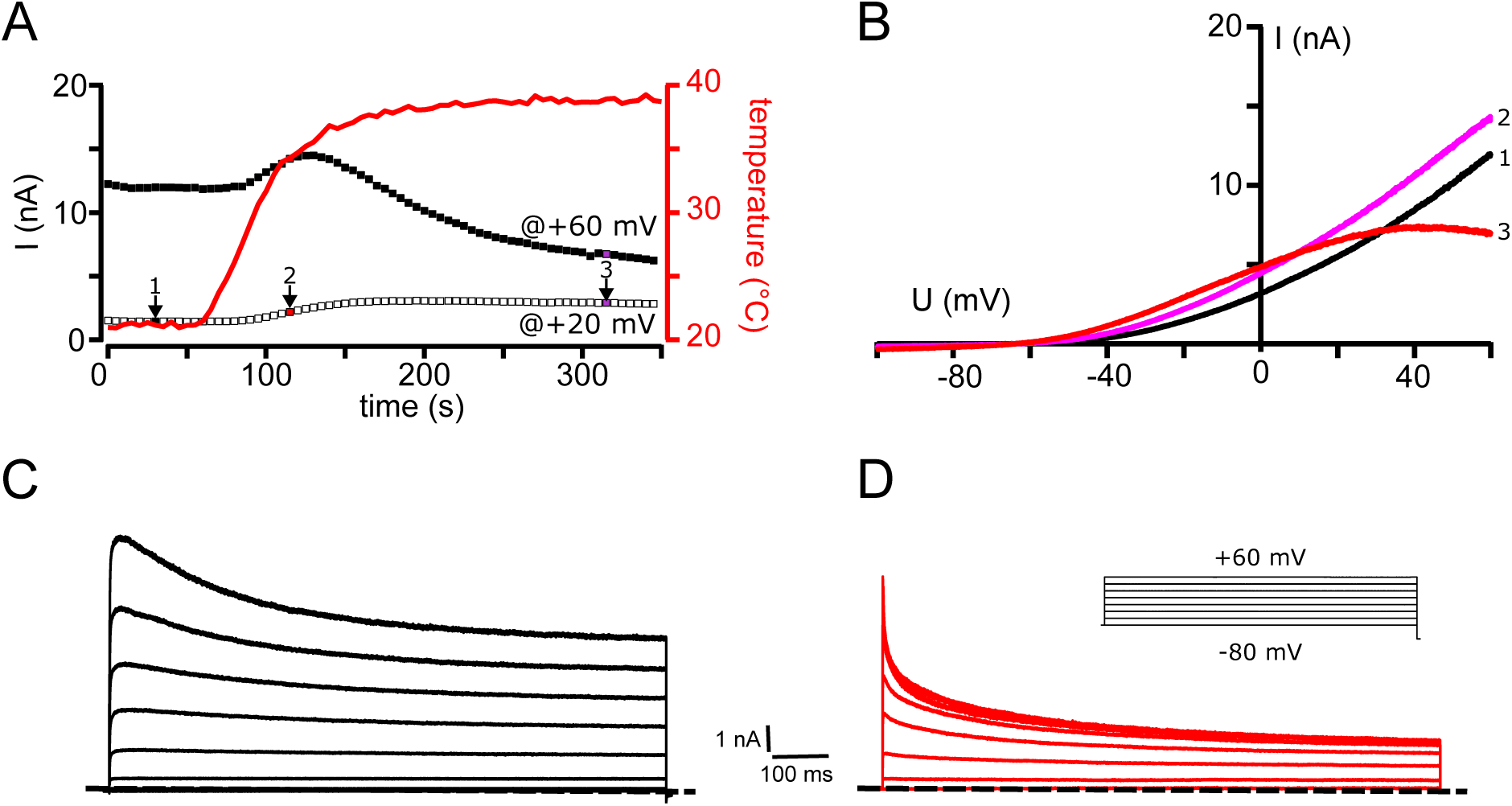
Temperature sensitivity of TWIK -1 channels. **A** Current response of TWIK-1 channels expressed in HEK293 cells to temperature elevation, measured with a ramp protocol (-100 to +60 mV; with Rb^+^ instead of K^+^ in the pipette solution). Red line: monitored temperature in the bath (filled squares: current at +60 mV; open squares: Current at +20 mV). **B** Current traces recorded with the ramp protocol at the time points indicated in A (1: at RT; 2: directly after temperature elevation; 3: with sustained temperature stimulation). **C** and **D** Current responses to families of rectangle pulses from -80 to +60 mV measured at RT (**C**) and at 37 °C (**D**), respectively.

**Figure S4.**
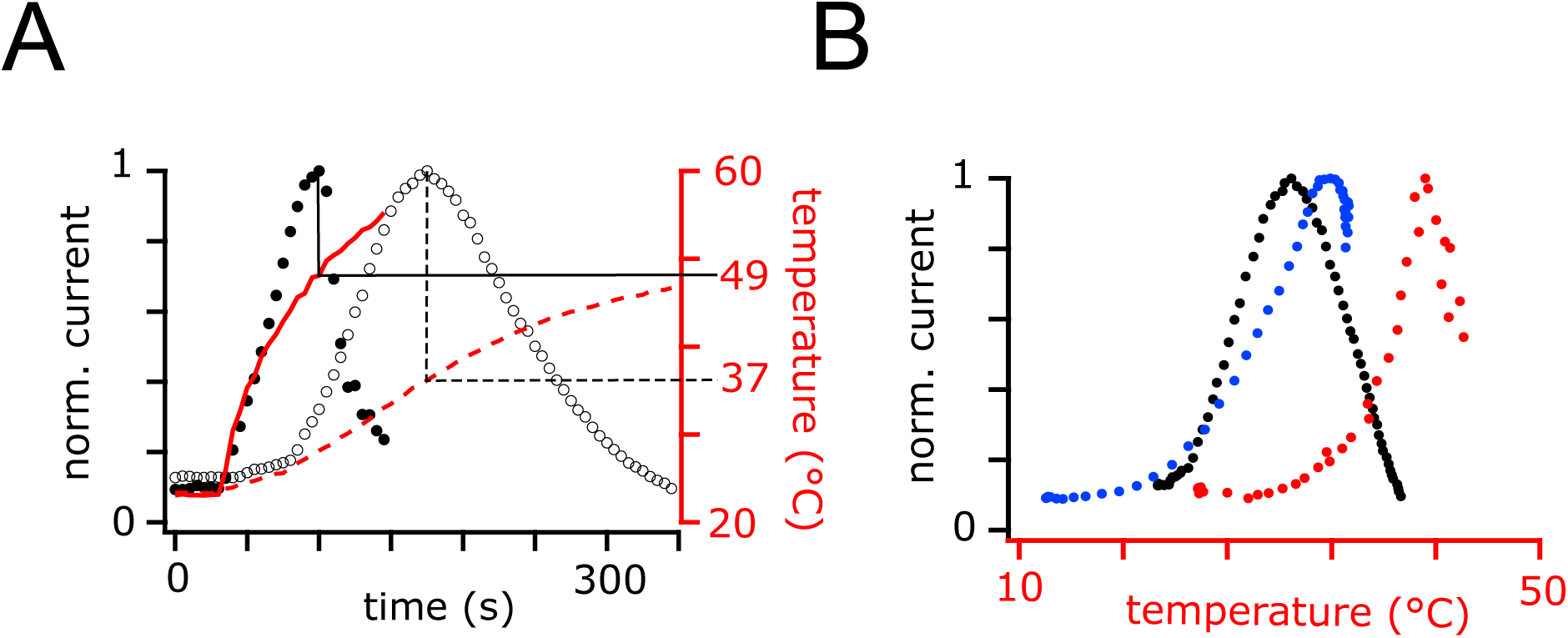
Temperature activation of TREK -1 channels in different recording setups. **A** The temperature at the current maximum depends on the temperature rise time. Faster temperature rise times result in maximal currents at higher temperature. Currents were measured with a ramp protocol and the currents at +60 mV were plotted against time. The currents were normalized to the maximal currents. Open circles/dotted lines mark currents elicited with a temperature rise time of 0.1 °C s^-1^. Filled circles/straight lines show currents elicited with a temperature rise time of 0.25 °C s^-1^. **B** Current changes as a function of the actual temperature for the measurements shown in A (black circles: temperature rise time 0.1 °C s^-1^; red circles: temperature rise time 0.25 °C s^-1^; blue circles: temperature rise time 0.1 °C s^-1^ with cooling prior to temperature elevation.

## References

Andharia, N. et al. (2017) ‘Involvement of intracellular transport in TREK-1c current run-up in 293T cells’, Channels, 11(3), pp. 224–235. Available at: 10.1080/19336950.2017.1279368.

Arrigoni, C. et al. (2016) ‘Unfolding of a Temperature-Sensitive Domain Controls Voltage-Gated Channel Activation’, Cell, 164(5), pp. 922–936. Available at: 10.1016/j.cell.2016.02.001.

Arrigoni, C. and Minor, D.L. (2018) ‘Global versus local mechanisms of temperature sensing in ion channels’, Pflugers Archiv: European Journal of Physiology, 470(5), pp. 733–744. Available at: 10.1007/s00424-017-2102-z.

Baas, P.W. et al. (2016) ‘Stability properties of neuronal microtubules’, *Cytoskeleton (Hoboken*, N.J.), 73(9), pp. 442–460. Available at: 10.1002/cm.21286.

Bagriantsev, S.N., Clark, K.A. and Minor, D.L. (2012) ‘Metabolic and thermal stimuli control K(2P)2.1 (TREK-1) through modular sensory and gating domains.’, The EMBO Journal, 31(15), pp. 3297–308. Available at: 10.1038/emboj.2012.171.

Bautista, D.M. et al. (2007) ‘The menthol receptor TRPM8 is the principal detector of environmental cold’, Nature, 448(7150), pp. 204–208. Available at: 10.1038/nature05910.

Bhave, G. et al. (2002) ‘cAMP-dependent protein kinase regulates desensitization of the capsaicin receptor (VR1) by direct phosphorylation’, Neuron, 35(4), pp. 721–731. Available at: 10.1016/s0896-6273(02)00802-4.

Brauchi, S. et al. (2006) ‘A hot-sensing cold receptor: C-terminal domain determines thermosensation in transient receptor potential channels’, The Journal of Neuroscience: The Official Journal of the Society for Neuroscience, 26(18), pp. 4835–4840. Available at: 10.1523/JNEUROSCI.5080-05.2006.

Brohawn, S.G., Su, Z. and MacKinnon, R. (2014) ‘Mechanosensitivity is mediated directly by the lipid membrane in TRAAK and TREK1 K+ channels.’, Proceedings of the National Academy of Sciences of the United States of America, 111(9), pp. 3614–9. Available at: 10.1073/pnas.1320768111.

Cao, E. et al. (2013) ‘TRPV1 channels are intrinsically heat sensitive and negatively regulated by phosphoinositide lipids’, Neuron, 77(4), pp. 667–679. Available at: 10.1016/j.neuron.2012.12.016.

Caterina, M.J. et al. (2000) ‘Impaired nociception and pain sensation in mice lacking the capsaicin receptor’, *Science (New York*, N.Y.), 288(5464), pp. 306–313. Available at: 10.1126/science.288.5464.306.

Chemin, J. et al. (2005) ‘Lysophosphatidic acid-operated K+ channels.’, The Journal of Biological Chemistry, 280(6), pp. 4415–21. Available at: 10.1074/jbc.M408246200.

Cho, H. et al. (2012) ‘The calcium-activated chloride channel anoctamin 1 acts as a heat sensor in nociceptive neurons’, Nature Neuroscience, 15(7), pp. 1015–1021. Available at: 10.1038/nn.3111.

Chowdhury, S., Jarecki, B.W. and Chanda, B. (2014) ‘A Molecular Framework for Temperature-Dependent Gating of Ion Channels’, Cell, 158(5), pp. 1148–1158. Available at: 10.1016/j.cell.2014.07.026.

Clapham, D.E. and Miller, C. (2011) ‘A thermodynamic framework for understanding temperature sensing by transient receptor potential (TRP) channels’, Proceedings of the National Academy of Sciences of the United States of America, 108(49), pp. 19492–19497. Available at: 10.1073/pnas.1117485108.

Colburn, R.W. et al. (2007) ‘Attenuated Cold Sensitivity in TRPM8 Null Mice’, Neuron, 54(3), pp. 379–386. Available at: 10.1016/j.neuron.2007.04.017.

Cui, Y. et al. (2012) ‘Selective disruption of high sensitivity heat activation but not capsaicin activation of TRPV1 channels by pore turret mutations’, The Journal of General Physiology, 139(4), pp. 273–283. Available at: 10.1085/jgp.201110724.

Davis, J.B. et al. (2000) ‘Vanilloid receptor-1 is essential for inflammatory thermal hyperalgesia’, Nature, 405(6783), pp. 183–187. Available at: 10.1038/35012076.

DeGiosio, R.A. et al. (2022) ‘More than a marker: potential pathogenic functions of MAP2’, Frontiers in Molecular Neuroscience, 15, p. 974890. Available at: 10.3389/fnmol.2022.974890.

Dehmelt, L. and Halpain, S. (2004) ‘The MAP2/Tau family of microtubule-associated proteins’, Genome Biology, 6(1), p. 204. Available at: 10.1186/gb-2004-6-1-204.

Delphin, C. et al. (2012) ‘MAP6-F is a temperature sensor that directly binds to and protects microtubules from cold-induced depolymerization’, The Journal of Biological Chemistry, 287(42), pp. 35127–35138. Available at: 10.1074/jbc.M112.398339.

Dhaka, A. et al. (2007) ‘TRPM8 is required for cold sensation in mice’, Neuron, 54(3), pp. 371–378. Available at: 10.1016/j.neuron.2007.02.024.

Dong, Y.Y. et al. (2015) ‘K2P channel gating mechanisms revealed by structures of TREK-2 and a complex with Prozac.’, Science, 347(6227), pp. 1256–1259. Available at: 10.1126/science.1261512.

Enyedi, P. and Czirják, G. (2010) ‘Molecular background of leak K+ currents: two-pore domain potassium channels.’, Physiological Reviews, 90(2), pp. 559–605. Available at: 10.1152/physrev.00029.2009.

Fink, M. et al. (1996) ‘Cloning, functional expression and brain localization of a novel unconventional outward rectifier K+ channel.’, The EMBO Journal, 15(24), pp. 6854–62.

Fink, M. et al. (1998) ‘A neuronal two P domain K+ channel stimulated by arachidonic acid and polyunsaturated fatty acids.’, The EMBO Journal, 17(12), pp. 3297–308. Available at: 10.1093/emboj/17.12.3297.

Fischer, M.J.M. and Reeh, P.W. (2007) ‘Sensitization to heat through G-protein-coupled receptor pathways in the isolated sciatic mouse nerve’, The European Journal of Neuroscience, 25(12), pp. 3570–3575. Available at: 10.1111/j.1460-9568.2007.05582.x.

Grandl, J. et al. (2008) ‘Pore region of TRPV3 ion channel is specifically required for heat activation’, Nature Neuroscience, 11(9), pp. 1007–1013. Available at: 10.1038/nn.2169.

Grandl, J. et al. (2010) ‘Temperature-induced opening of TRPV1 ion channel is stabilized by the pore domain’, Nature Neuroscience, 13(6), pp. 708–714. Available at: 10.1038/nn.2552.

Hille, B. (2001) Ion Channels of Excitable Membranes. 3rd edn. Edited by B. Hille. Sunderland, MA: Sinauer Associates.

Honoré, E. (2007) ‘The neuronal background K2P channels: focus on TREK1.’, Nature reviews. Neuroscience, 8(4), pp. 251–61. Available at: 10.1038/nrn2117.

Hucho, T. and Levine, J.D. (2007) ‘Signaling Pathways in Sensitization: Toward a Nociceptor Cell Biology’, Neuron, 55(3), pp. 365–376. Available at: 10.1016/j.neuron.2007.07.008.

Jabba, S. et al. (2014) ‘Directionality of temperature activation in mouse TRPA1 ion channel can be inverted by single-point mutations in ankyrin repeat six’, Neuron, 82(5), pp. 1017–1031. Available at: 10.1016/j.neuron.2014.04.016.

Kang, D., Choe, C. and Kim, D. (2005) ‘Thermosensitivity of the two-pore domain K+ channels TREK-2 and TRAAK.’, The Journal of Physiology, 564(Pt 1), pp. 103–116. Available at: 10.1113/jphysiol.2004.081059.

Kwon, D.H. et al. (2021) ‘Heat-dependent opening of TRPV1 in the presence of capsaicin’, Nature structural & molecular biology, 28(7), p. 554. Available at: 10.1038/s41594-021-00616-3.

Lamas, J.A., Rueda-Ruzafa, L. and Herrera-Pérez, S. (2019) ‘Ion Channels and Thermosensitivity: TRP, TREK, or Both?’, International Journal of Molecular Sciences, 20(10), p. 2371. Available at: 10.3390/ijms20102371.

Maingret, F. et al. (1999) ‘Mechano- or Acid Stimulation, Two Interactive Modes of Activation of the TREK-1 Potassium Channel’, The Journal of Biological Chemistry, 274(38), pp. 26691–26696. Available at: 10.1074/jbc.274.38.26691.

Maingret, F. et al. (2000) ‘TREK-1 is a heat-activated background K(+) channel.’, The EMBO Journal, 19(11), pp. 2483–91. Available at: 10.1093/emboj/19.11.2483.

Marsh, B. et al. (2012) ‘Leak K+ channel mRNAs in dorsal root ganglia: Relation to inflammation and spontaneous pain behaviour’, Molecular and Cellular Neuroscience, 49(3), pp. 375–386. Available at: 10.1016/j.mcn.2012.01.002.

McClenaghan, C. et al. (2016) ‘Polymodal activation of the TREK-2 K2P channel produces structurally distinct open states.’, The Journal of General Physiology, 147(6), pp. 497–505. Available at: 10.1085/jgp.201611601.

Melková, K. et al. (2019) ‘Structure and Functions of Microtubule Associated Proteins Tau and MAP2c: Similarities and Differences’, Biomolecules, 9(3), p. 105. Available at: 10.3390/biom9030105.

Moparthi, L. et al. (2022) ‘The human TRPA1 intrinsic cold and heat sensitivity involves separate channel structures beyond the N-ARD domain’, Nature Communications, 13(1), p. 6113. Available at: 10.1038/s41467-022-33876-8.

Morrison, S.F., Nakamura, K. and Madden, C.J. (2008) ‘Central control of thermogenesis in mammals’, Experimental Physiology, 93(7), pp. 773–797. Available at: 10.1113/expphysiol.2007.041848.

Murbartián, J. et al. (2005) ‘Sequential phosphorylation mediates receptor- and kinase-induced inhibition of TREK-1 background potassium channels.’, The Journal of Biological Chemistry, 280(34), pp. 30175–84. Available at: 10.1074/jbc.M503862200.

Nadezhdin, K.D. et al. (2021) ‘Structural mechanism of heat-induced opening of a temperature-sensitive TRP channel’, Nature Structural & Molecular Biology, 28(7), pp. 564–572. Available at: 10.1038/s41594-021-00615-4.

Nakamura, K. (2024) ‘Central Mechanisms of Thermoregulation and Fever in Mammals’, Advances in Experimental Medicine and Biology, 1461, pp. 141–159. Available at: 10.1007/978-981-97-4584-5_10.

Noël, J. et al. (2009) ‘The mechano-activated K+ channels TRAAK and TREK-1 control both warm and cold perception.’, The EMBO Journal, 28(9), pp. 1308–18. Available at: 10.1038/emboj.2009.57.

Obar, R.A. et al. (1989) ‘The RII subunit of camp-dependent protein kinase binds to a common amino-terminal domain in microtubule-associated proteins 2A, 2B, and 2C’, Neuron, 3(5), pp. 639–645. Available at: 10.1016/0896-6273(89)90274-2.

Palkar, R., Lippoldt, E.K. and McKemy, D.D. (2015) ‘The molecular and cellular basis of thermosensation in mammals’, Current opinion in neurobiology, 34, p. 14. Available at: 10.1016/j.conb.2015.01.010.

Patel, A.J. et al. (1998) ‘A mammalian two pore domain mechano-gated S-like K+ channel.’, The EMBO Journal, 17(15), pp. 4283–90. Available at: 10.1093/emboj/17.15.4283.

Patel, A.J. et al. (2000) ‘TWIK-2, an Inactivating 2P Domain K+ Channel’, Journal of Biological Chemistry, 275(37), pp. 28722–28730. Available at: 10.1074/jbc.M003755200.

Pereira, V. et al. (2014) ‘Role of the TREK2 potassium channel in cold and warm thermosensation and in pain perception’, Pain, 155(12), pp. 2534–2544. Available at: 10.1016/j.pain.2014.09.013.

Ranels, H.J. and Griffin, J.D. (2003) ‘The effects of prostaglandin E2 on the firing rate activity of thermosensitive and temperature insensitive neurons in the ventromedial preoptic area of the rat hypothalamus’, Brain Research, 964(1), pp. 42–50. Available at: 10.1016/S0006-8993(02)04063-5.

Ranels, H.J. and Griffin, J.D. (2005) ‘Effects of prostaglandin E2 on the electrical properties of thermally classified neurons in the ventromedial preoptic area of the rat hypothalamus’, BMC neuroscience, 6, p. 14. Available at: 10.1186/1471-2202-6-14.

Sandoz, G. et al. (2006) ‘AKAP150, a switch to convert mechano-, pH- and arachidonic acid-sensitive TREK K(+) channels into open leak channels.’, The EMBO Journal, 25(24), pp. 5864–72. Available at: 10.1038/sj.emboj.7601437.

Sandoz, G. et al. (2008) ‘Mtap2 is a constituent of the protein network that regulates twik-related K+ channel expression and trafficking.’, The Journal of Neuroscience: The Official Journal of the Society for Neuroscience, 28(34), pp. 8545–52. Available at: 10.1523/JNEUROSCI.1962-08.2008.

Sengupta, P. and Garrity, P. (2013) ‘Sensing temperature’, Current Biology, 23(8), pp. R304–R307. Available at: 10.1016/j.cub.2013.03.009.

Singh, A.K. et al. (2019) ‘Structural basis of temperature sensation by the TRP channel TRPV3’, Nature Structural & Molecular Biology, 26(11), pp. 994–998. Available at: 10.1038/s41594-019-0318-7.

Talley, E.M. et al. (2001) ‘Cns distribution of members of the two-pore-domain (KCNK) potassium channel family.’, The Journal of Neuroscience: The Official Journal of the Society for Neuroscience, 21(19), pp. 7491–505. Available at: 10.1523/JNEUROSCI.21-19-07491.2001.

Tan, C.L. and Knight, Z.A. (2018) ‘Regulation of body temperature by the nervous system’, Neuron, 98(1), p. 31. Available at: 10.1016/j.neuron.2018.02.022.

Türkaydin, B. et al. (2024) ‘Atomistic mechanism of coupling between cytosolic sensor domain and selectivity filter in TREK K2P channels’, Nature Communications, 15(1), p. 4628. Available at: 10.1038/s41467-024-48823-y.

Viatchenko-Karpinski, V., Ling, J. and Gu, J.G. (2018) ‘Characterization of temperature-sensitive leak K+ currents and expression of TRAAK, TREK-1, and TREK2 channels in dorsal root ganglion neurons of rats’, Molecular Brain, 11(1), p. 40. Available at: 10.1186/s13041-018-0384-5.

Vriens, J., Nilius, B. and Voets, T. (2014) ‘Peripheral thermosensation in mammals’, Nature Reviews. Neuroscience, 15(9), pp. 573–589. Available at: 10.1038/nrn3784.

Watanabe, H. et al. (2002) ‘Heat-evoked activation of TRPV4 channels in a HEK293 cell expression system and in native mouse aorta endothelial cells’, The Journal of Biological Chemistry, 277(49), pp. 47044–47051. Available at: 10.1074/jbc.M208277200.

Wechselberger, M. et al. (2006) ‘Ionic channels and conductance-based models for hypothalamic neuronal thermosensitivity’, *American Journal of Physiology-Regulatory*, Integrative and Comparative Physiology, 291(3), pp. R518–R529. Available at: 10.1152/ajpregu.00039.2006.

Xiao, B. et al. (2011) ‘Temperature-dependent STIM1 activation induces Ca2+ influx and modulates gene expression’, Nature chemical biology, 7(6), p. 351. Available at: 10.1038/nchembio.558.

Yamamoto, Y., Hatakeyama, T. and Taniguchi, K. (2009) ‘Immunohistochemical colocalization of TREK-1, TREK-2 and TRAAK with TRP channels in the trigeminal ganglion cells’, Neuroscience Letters, 454(2), pp. 129–133. Available at: 10.1016/j.neulet.2009.02.069.

Yang, F. et al. (2010) ‘Thermosensitive TRP channel pore turret is part of the temperature activation pathway’, Proceedings of the National Academy of Sciences, 107(15), pp. 7083–7088. Available at: 10.1073/pnas.1000357107.

Yao, J., Liu, B. and Qin, F. (2011) ‘Modular thermal sensors in temperature-gated transient receptor potential (TRP) channels’, Proceedings of the National Academy of Sciences of the United States of America, 108(27), pp. 11109–11114. Available at: 10.1073/pnas.1105196108.

Yeh, F., Jara-Oseguera, A. and Aldrich, R.W. (2023) ‘Implications of a temperature-dependent heat capacity for temperature-gated ion channels’, Proceedings of the National Academy of Sciences of the United States of America, 120(24), p. e2301528120. Available at: 10.1073/pnas.2301528120.

Zakharian, E., Cao, C. and Rohacs, T. (2010) ‘Gating of transient receptor potential melastatin 8 (TRPM8) channels activated by cold and chemical agonists in planar lipid bilayers’, The Journal of Neuroscience: The Official Journal of the Society for Neuroscience, 30(37), pp. 12526–12534. Available at: 10.1523/JNEUROSCI.3189-10.2010.

Zhang, F. et al. (2018) ‘Heat activation is intrinsic to the pore domain of TRPV1’, Proceedings of the National Academy of Sciences of the United States of America, 115(2), pp. E317–E324. Available at: 10.1073/pnas.1717192115.

Zimmermann, K. et al. (2007) ‘Sensory neuron sodium channel Nav1.8 is essential for pain at low temperatures’, Nature, 447(7146), pp. 855–858. Available at: 10.1038/nature05880.

